## Supplemental Table 1 for "Multiple dynamic interactions from basal ganglia direct and indirect pathways mediate action selection"

**Key resources table**

| REAGENT or RESOURCE | SOURCE | IDENTIFIER |
| --- | --- | --- |
| Bacterial and virus strains | | |
| AAV9-FLEX-DTR-GFP | Salk GT3 Core | N/A |
| AAV5-EF1a-DIO-hChR2(H134R)-mCherry | University of North Carolina  Vector Core | N/A |
| AAV9-EF1a-DIO-hChR2(H134R)-eYFP | University of Pennsylvania  Vector Core | Cat# AV-9-20298P |
| AAV5-EF1a-DIO-eNpHR3.0-eYFP | University of North Carolina  Vector Core | N/A |
| Chemicals, peptides, and recombinant proteins | | |
| Muscimol, GABAA receptor agonist | Sigma-Aldrich | Cat# M1523 |
| Diphtheria toxin | List Biological Labs | Part# 150 |
| Experimental models: Organisms/strains | | |
| Mouse: C57BL/6 | Envigo/Harlan | Code: 044 |
| Mouse: NR1f/f (B6.129S4-Grin1tm2Stl/J) | Jackson Laboratory | Stock# 005246 |
| Mouse: Ai32 (B6;129S-Gt(ROSA)26Sortm32(CAG-COP4*H134R/EYFP)Hze/J) | Jackson Laboratory | Stock# 012569 |
| Mouse: D1-cre (B6.FVB(Cg)-Tg(Drd1a-cre)EY217Gsat/Mmucd) | MMRRC | RRID: MMRRC_034258-UCD |
| Mouse: A2a-cre (B6.FVB(Cg)-Tg(Adora2a-cre)KG139Gsat/Mmucd) | MMRRC | RRID: MMRRC_036158-UCD |
| Software and algorithms | | |
| GraphPad Prism | GraphPad Software | Version 7.03;  https://www.graphpad.com/  scientific-software/prism/ |
| MATLAB | MathWorks | R2013a;  https://www.mathworks.com/products/matlab.html |
| Med-PC | Med Associates | Cat# SOF-735;  http://www.med-associates.com/  product/med-pc-iv-software/ |
| Offline Sorter | Plexon | Version 3.3.3;  https://plexon.com/products/offline-sorter/ |
| OmniPlex | Plexon | Version 1.4.5;  https://plexon.com/products/omniplex-software/ |
| EthoVision | Noldus | Version 8.5 |
| Other | | |
| Med Associates operant chamber | Med Associates | Cat# MED-307W-D1 |
| Electrode Array | Innovative Neurophysiology | N/A |
| 473 nm laser | LaserGlow Technologies | N/A |
| 532 nm laser | LaserGlow Technologies | N/A |
